# Self-Supervised Maize Kernel Classification and Segmentation for Embryo Identification

**DOI:** 10.1101/2022.11.25.517990

**Authors:** David Dong, Koushik Nagasubramanian, Ruidong Wang, Ursula K Frei, Talukder Z Jubery, Thomas Lübberstedt, Baskar Ganapathysubramanian

## Abstract

Computer vision and deep learning (DL) techniques have succeeded in a wide range of diverse fields. Recently, these techniques have been successfully deployed in plant science applications to address food security, productivity, and environmental sustainability problems for a growing global population. However, training these DL models often necessitates the large-scale manual annotation of data which frequently becomes a tedious and time-and-resource-intensive process. Recent advances in self-supervised learning (SSL) methods have proven instrumental in overcoming these obstacles, using purely unlabeled datasets to pre-train DL models. Here, we implement the popular self-supervised contrastive learning methods of NNCLR (Nearest neighbor Contrastive Learning of visual Representations) and SimCLR (Simple framework for Contrastive Learning of visual Representations) for the classification of spatial orientation and segmentation of embryos of maize kernels. Maize kernels are imaged using a commercial high-throughput imaging system. This image data is often used in multiple downstream applications across both production and breeding applications, for instance, sorting for oil content based on segmenting and quantifying the scutellum’s size and for classifying haploid and diploid kernels. We show that in both classification and segmentation problems, SSL techniques outperform their purely supervised transfer learning-based counterparts and are significantly more annotation efficient. Additionally, we show that a single SSL pre-trained model can be efficiently finetuned for both classification and segmentation, indicating good transferability across multiple downstream applications. Segmentation models with SSL-pretrained backbones produce DICE similarity coefficients of 0.81, higher than the 0.78 and 0.73 of those with ImageNet-pretrained and randomly initialized backbones, respectively. We observe that finetuning classification and segmentation models on as little as 1% annotation produces competitive results. These results show SSL provides a meaningful step forward in data efficiency with agricultural deep learning and computer vision.

## 1. INTRODUCTION

Deep learning (DL) for computer vision applications has recently become a boon to innovations in agricultural efficiency. These methods have transformed how we extract various agronomically relevant plant traits under laboratory and field conditions [1-4]. Automatically and rapidly extracting plant traits can be a game-changer in terms of reducing food costs and improving production efficiencies, improving sustainability by reducing waste, and providing a better understanding of adapting crops for climate change. Deep learning methods have been used in various agricultural applications to identify, classify, quantify, and predict traits [5-9]. With the availability of high-throughput data acquisition tools that produce large amounts of good-quality data, the major bottleneck in deploying DL-based computer vision tools is the need for large amounts of labeled data to train these DL models. Data annotation or labeling is the main development barrier to building high-quality DL models, especially since labeling the raw data often requires domain experts to annotate images. Data annotation by an expert with domain-specific knowledge is a tedious and expensive task. The DL community is exploring various strategies to break this dependency on a large quantity of annotated data to train DL models in a label-efficient manner, including approaches like active learning [10], transfer learning [11], weakly supervised learning [12, 13] and the more recent advances in self-supervised learning [14, 15]. In this work, we focus on deploying self-supervised learning approaches to the problem of characterizing maize kernels that are imaged in a commercial high-throughput seed imaging system (Qsorter technologies [16]). We consider two vision tasks – first, identify if the maize kernels are correctly oriented for downstream analysis (a classification task), and second, segment out the kernel scutellum from the correctly oriented seeds (a segmentation task).

The ability to accurately and efficiently segment maize kernel scutellum has significant utility for both production and breeding application. Maize oil (corn oil) is extracted from corn kernels through milling [17]. Milling processes are integrated into the production of corn starch, sugar, syrup, alcohol, and byproducts like gluten feed, along with corn oil. Of the 1.1 billion metric tons of corn produced annually around the world, over 3.5 million are used for oil production [18, 19]. Almost all oil is found in the embryo of the kernel [17]. The ability to sort seeds for embryo/scutellum size is a significant value addition. Similarly, the non-destructive sorting of single seeds based on oil content (OC) has been shown to be useful for early-generation screening to improve the efficiency of breeding [20, 21] and for haploid selection in an oil-inducer-based doubled haploid breeding program [22, 23]. Over the past few years, nuclear magnetic resonance (NMR) [24, 25], fluorescence imaging [26], near-infrared (NIR) reflectance spectroscopy [27-30], hyperspectral imaging [31], and line-scan Raman hyperspectral imaging [32] have been developed to measure or predict oil content. However, these methods and tools are expensive. On the other hand, sorting based on NIR reflectance is less costly, has been around for a long time [33, 34], and has worked well to predict protein and starch content. However, using those tools to measure OC is not easy because the position of the embryo/scutellum to the camera/light source [35] strongly affects OC measurements of single seeds, which leads to significant standard errors. Several currently available NIR spectra-based high throughput single seed sorting devices capture RGB images of the seed along with the NIR spectrum [16, 36]. These images can be used to identify the correctly orientated seed and quantify the relative size of the embryo to the seed, which, coupled with the NIR spectrum, could be used to improve the prediction of OC.

This work aims to design an end-to-end DL framework that classifies kernels and segments the embryo. Accurately performing these steps will allow us to, in the future, predict corn OC with high accuracy. **Figure 1** illustrates this pipeline. A challenge in accomplishing this goal is that DL techniques often rely on having access to large datasets of annotated images for successful training results. This problem motivates our approach of using self-supervised contrastive. The self-supervised pretraining procedure automatically uses unlabeled data to generate pretrained labels [37]. It does so by solving a pretext task suited for learning representations, which in computer vision typically consists of learning invariance to image augmentations like rotation and color transforms, producing feature representations that ideally can be easily adapted for use in a downstream task. After obtaining this pre-trained model, we apply standard DL to finetune the model with a smaller labeled dataset. The smaller labeled dataset is used to reduce the effect of possible inaccuracies in the pseudo-labels from the self-supervised task [38]. The orientation of corn kernels must maintain consistency between measurements and be oriented to fully display the embryo. The goal of the segmentation problem is then to identify the embryo amidst the background and the rest of each kernel.

**Figure 1.**
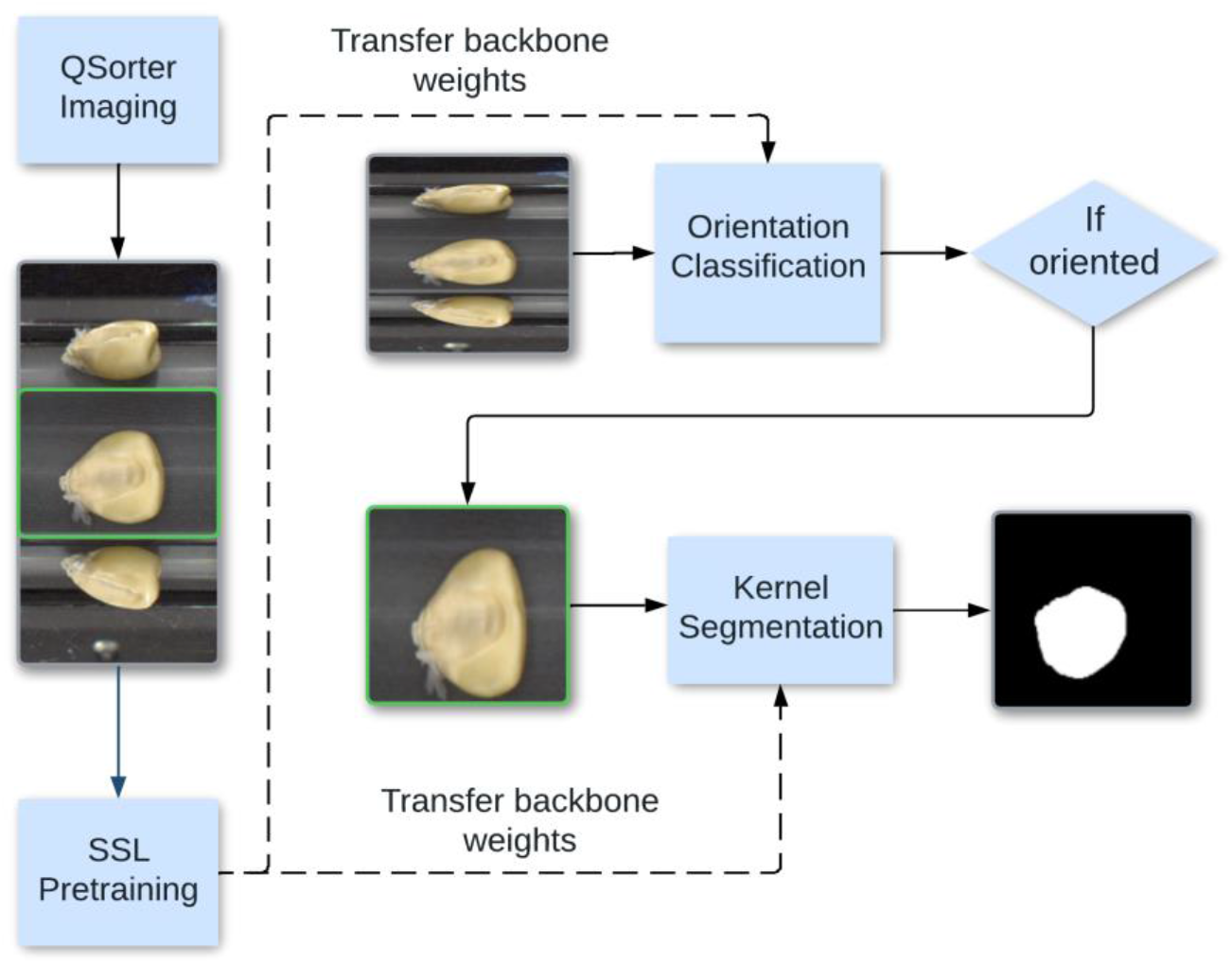
End-to-end pipeline for corn kernel classification and segmentation. The curved arrow shows the middle pane being processed for the segmentation task. We show that the classification and segmentation models perform bets with self-supervised weights.

Our contributions in this paper are 1) the creation of an end-to-end DL pipeline for kernel classification and segmentation, facilitating downstream applications in OC prediction, 2) to assess capabilities of self-supervised learning regarding annotation efficiency, and 3) illustrating the ability of self-supervised pretraining to create models that can be finetuned for diverse downstream applications. Beyond the direct application of the classification and segmentation capabilities of the learned representations, using self-supervised techniques, in general, could accelerate the development of computer vision techniques for ag applications, skipping several stages of arduous and time-consuming data collection.

## 2. MATERIALS AND METHODS

### 2.1 Dataset

#### 2.1.1 Dataset for classification by imaging orientation

The classification dataset consists of 44,286 RGB 492-pixel by 240-pixel images of maize kernels of various accessions taken using the RGB imaging tools of QSorter. Of these, 2697 were manually labeled into two classes: “oriented” and “non-oriented.” Kernels that belong to the “oriented” class were deemed appropriate for calculating internal OC within the embryo/germ center of corn kernels. This determination was based on the requirement that the visible embryo is parallel to the camera’s plane.

In a typical downstream application, this visual information provided by image segmentation would be combined with data from the hyperspectral imaging sensor provided by QSorter, but with such a sensor having its field of view limited to only the middle pane. However, the other two panes still provide useful visual information for our classification models since the determination of the orientation of any particular kernel is not limited to only the frontal view of the kernel. **Figure 2**a shows oriented kernels, noting the lighter portion visible in each middle pane, which is the corn embryo’s visible part. **Figure 2**b shows non-oriented kernels in which the embryos are not visible or only partially visible.

**Figure 2:**
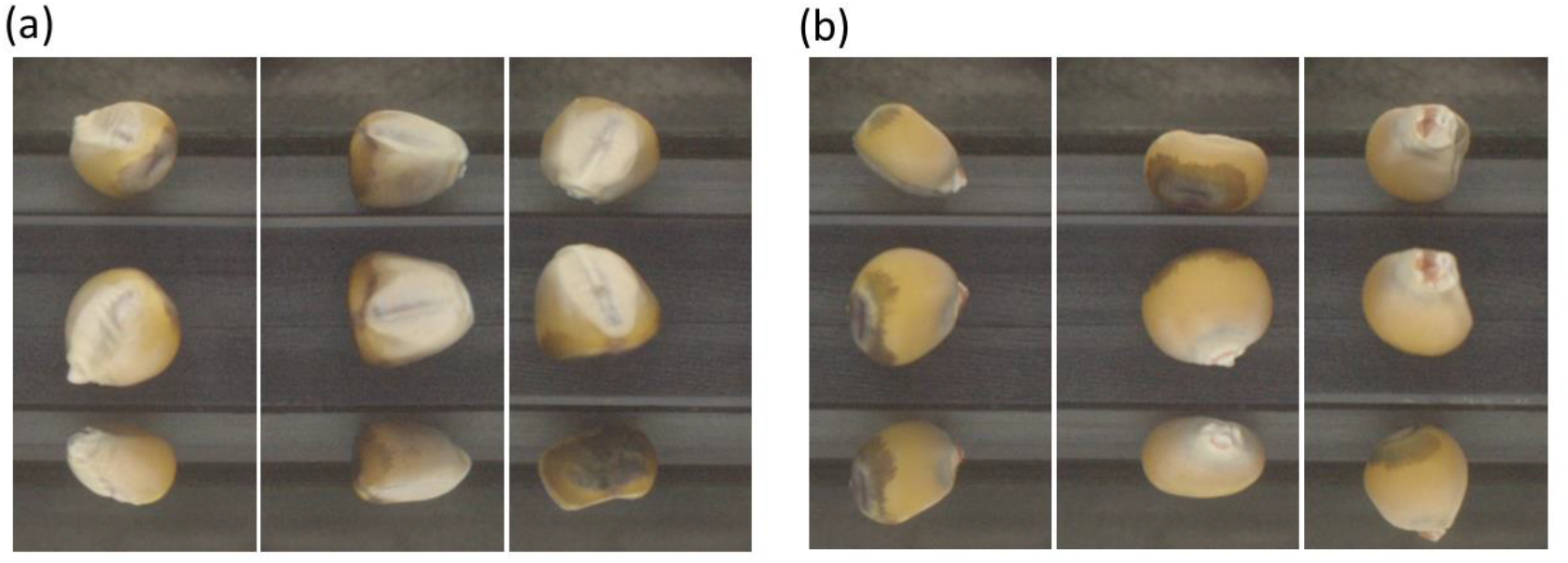
Images classified as “oriented” with the embryo visible (a) and “non-oriented” with the embryo not visible (b).

#### 2.2.2 Dataset for embryo segmentation

The embryo segmentation dataset consists of only 401 RGB images of corn kernels, taken from the same source of QSorter images as in the classification dataset above, along with their respective binary masks. Thus, the 2D image shapes were again 492 × 240. Segmentation (into the binary mask) distinguishes between the embryo and the rest of the background (including the non-embryo portion of the kernel). **Figure 3** illustrates the segmentation annotation process for an RGB image and its mask. The three frames of each original (492, 240) dataset image were split into three individual images and downsampled to (128, 128). All completely negative masks and their respective RGB images were then removed.

**Figure 3.**
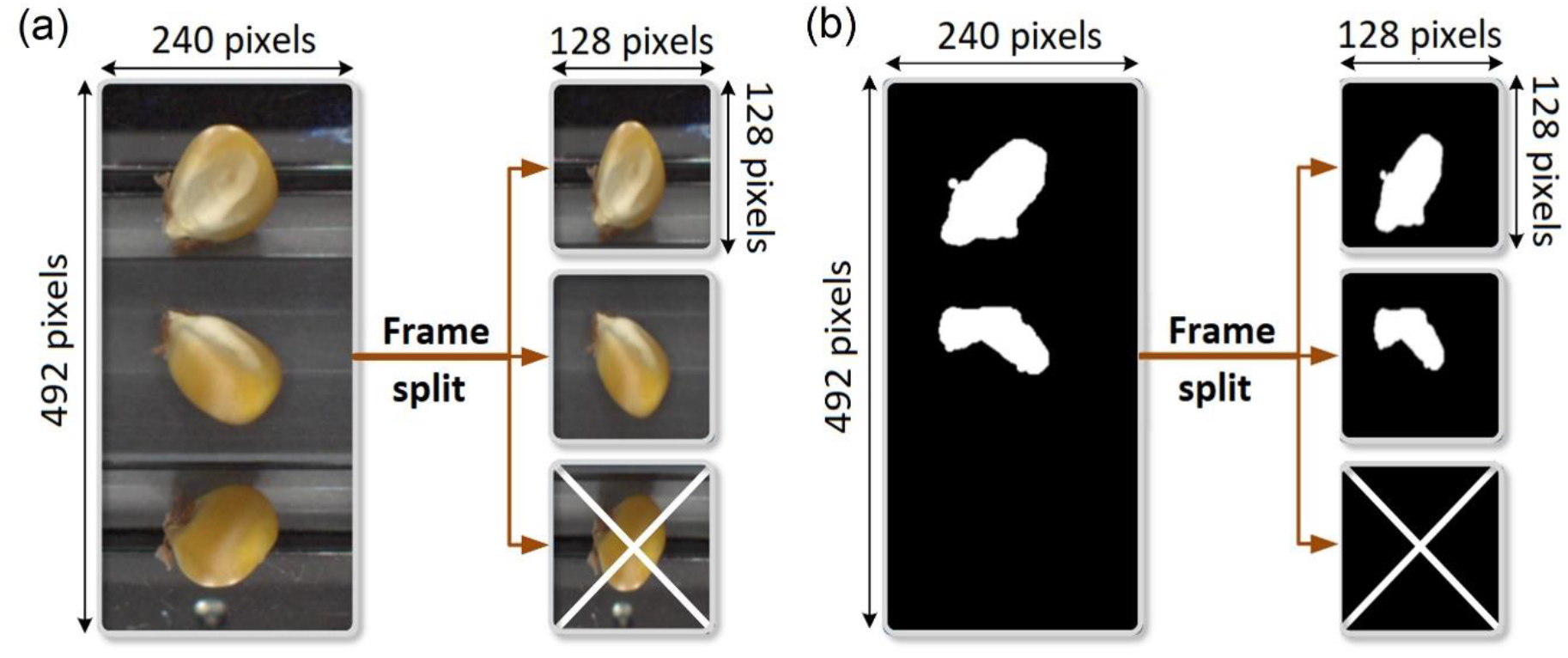
Preprocessing for segmentation consists of splitting each dataset image into three 128 × 128 images. Completely negative masks were excluded. (a) Preprocessing of RGB image. (b) Preprocessing of the mask.

### 2.2 SSL Pretraining

#### 2.2.1 Methods overview

The contrastive learning framework is a self-supervised learning method that maximizes the similarity between representations of an image and the augmented version of an image while minimizing the similarity between an image and other images [39]. The two models used for self-supervised pretraining were SimCLR (Simple Framework for Contrastive Learning of Visual Representations) [40] and NNCLR (Nearest-Neighbor Contrastive Learning of Visual Representations) [41]. **Figure 4** shows these two models superimposed on the same diagram.

**Figure 4.**
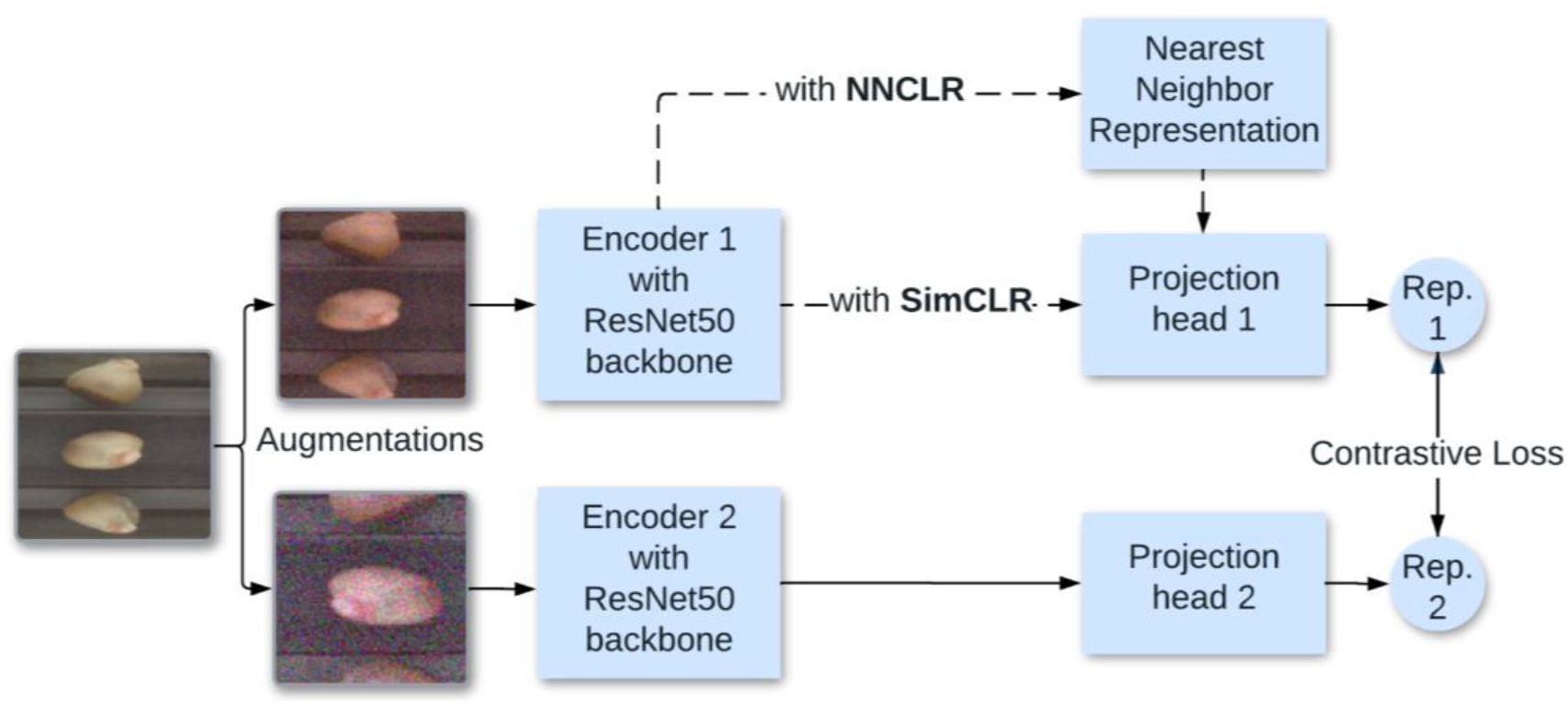
Superimposed diagram of NNCLR and SimCLR models. The two dashed arrows show the paths a tensor representation passes depending on the self-supervised model.

SimCLR trains a backbone used for downstream processes by considering the contrastive loss of the representations of two distinct augmentations of images extracted from any given batch. If the initial images are the same, the pair of representations is considered a positive pair for the final calculation, and if the views are augmentations of two distinct images in the batch, then it is considered a negative pair. The representations are created by taking each augmented view of the initial image along a path including two networks: a base encoder where the desired backbone resides and a final projection head to calculate the contrastive loss of the representation in a separate space. NNCLR is also a contrastive model but differs from SimCLR in that upon taking both views of a given image through an encoder; the nearest neighbor algorithm is used to sample dataset representations for one of the views from a subset of the initial dataset. These are treated as the analog of the positive pairs described in the SimCLR model. Negative pairs are then the nearest neighbors of distinct initial images. Both architectures use the same InfoNCE loss to maximize agreement, a loss function using categorical cross-entropy to maximize agreement with positive samples, commonly used in self-supervised learning [42]. To evaluate the performance of the pretrained models, a linear probe — separate from the non-linear projection head included in both models — was attached directly to the encoder and was weight-updated at each step. The backbone and probe were then extracted to calculate validation accuracy for model selection.

#### 2.2.2 Contrastive data augmentation

In many supervised image processing and computer vision tasks, data augmentation is used for the dual purposes of increasing the size of a labeled dataset through synthetic means and improving the diversity of a dataset. For purely supervised purposes, data augmentation can synthetically multiply the dataset’s size by altering existing data and increasing the diversity of data to generalize the training set better [43]. Contrastive learning uses heavier image augmentations than would normally be supplied to purely supervised training [44]. This is due to the reliance of contrastive learning on using augmentations as a model for learning invariance to “style” changes, while the “content” component of a representation remains invariant [45]. Thus, heavy stylistic changes should generally benefit the learned representations.

The data augmentations used for our pretraining process were derived from the recommended augmentations particular to SimCLR, consisting of random zoom, random flip, color jitter, and Gaussian noise. NNCLR is less dependent in its performance than SimCLR on the precise type and magnitude of data augmentations used in training; indeed, upon applying augmentations to NNCLR pretraining similar to the full set recommended for SimCLR produced only a 1.6% performance improvement when compared to using only random crop [41].

#### 2.2.3 Pretraining setup

Hyperparameter sweeping during pretraining consisted of the variation of the contrastive learning rate, the type of weight initialization applied to the ResNet50 backbone, and data augmentation strength. The learning rate was chosen between 1e-3 and 1e-4, coupling the contrastive learning rate with the classification learning rate of the linear probe. Weight initialization was chosen between ImageNet and random initialization. The data augmentation strength of each augmentation was varied together and explained below. Thus, eight runs were processed for each sweep, and each sweep was repeated three times to ensure precision.

### 2.3 Classification

#### 2.3.1 Data split

Of the 2697 images manually classified from the unlabeled dataset, there were 1300 oriented images and 1367 non-oriented images. Of the labeled images, 1697 were used for training, with an 800:897 class split in favor of non-oriented images. The rest were divided between validation and testing and were split evenly between the classes. So, 500 images were allocated to each set, with 250 images in each class. During pretraining, the images allocated to the validation and testing were separated from the unlabeled dataset used for contrastive learning, while the labeled training dataset was included, such that 43,286 out of the 44286 total images were used for unlabeled contrastive learning.

#### 2.3.2 Training setup

The training process was set up to facilitate comparison between different models after undergoing end-to-end finetuning. Only ResNet50 was used for the backbones, as is standard in self-supervised model evaluation and as was used in both the NNCLR and SimCLR original papers [40, 41, 46]. Two backbones for the end-to-end process were chosen from a pretraining sweep with the mentioned self-supervised contrastive architectures, and one backbone was initialized with ImageNet weights.

Data augmentation strength was defined separately for each particular augmentation depending on its configuration specifics: Random zoom acted by cropping to a single rectangle with its shape uniformly chosen between a maximum area of the initial 128×128 2D image shape and a minimum area of either 25% or 75% of the maximum area. Brightness and color transform was accomplished first by taking an identity matrix multiplied by the chosen brightness factor, then adding a matrix with uniformly chosen values selected between the jitter factor and its negative, and secondly by multiplying the original dataset image by this matrix. Brightness jitter increased the brightness of the image by either 50% or 75%, and the jitter factor was either 0.3 or 0.45. Gaussian noise was applied with a standard deviation of either 0.1 or 1.5. The only augmentation kept constant was random flip, constantly at 50% activation chance. Upon evaluation, the two chosen models from this pretraining sweep process—corresponding to the top-left-most two light-green boxes in **Figure 5**—were backbones pretrained by 1) NNCLR with random initialization at LR = 1E-3 and 2) SimCLR with ImageNet initialization at LR = 1E-3.

**Figure 5:**
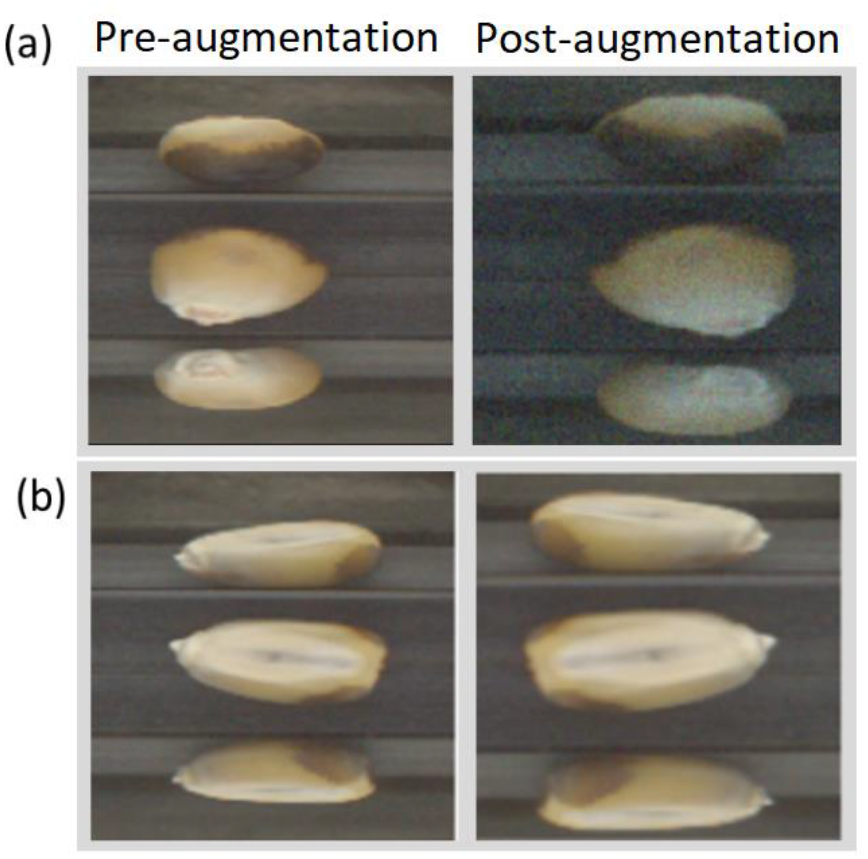
Two examples of a processed dataset image and its augmentation. (a) Random zoom, horizontal flip, brightness and color transform, and Gaussian noise. (b) Random zoom and horizontal flip.

#### 2.3.4 Feature extraction and finetuning

During training, separate trials were performed for each proportion of annotated data used in classification (1%, 10%, 25%, 100%). As in pretraining, each trial was repeated three times. With 1% and 10% data, a batch size of 4 was used; for 25% data, a batch size of 32 was used; and for 100% data, a batch size of 128 was used. During feature extraction, first, the ResNet-50 backbone from each initialization method was frozen to weight updates, upon which a trainable one-node classifier was constructed with sigmoid activation. Each classifier in every trial was trained for 300 epochs. In finetuning, the backbone was unfrozen, and the entire model was trained for 400 epochs. The same learning rate schedule was used in both phases at the fixed schedule of a 0.5 multiplier every 50 epochs. This process is illustrated in **Figure 6**.

**Figure 6.**
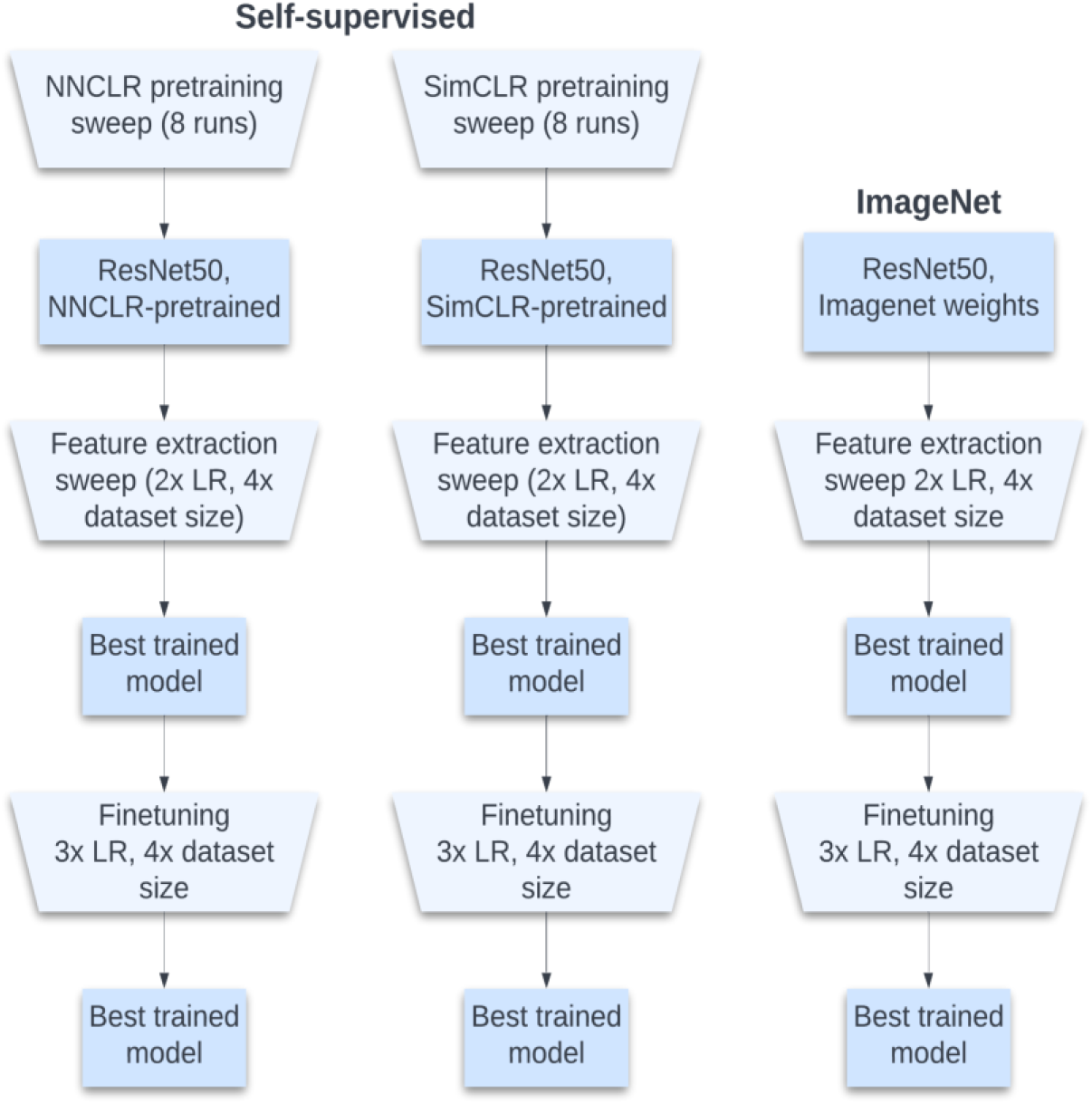
Training process. With each sweep over hyperparameters, the best model is chosen for the next round.

### 2.4 Segmentation

Semantic segmentation is a pixel-level classification problem where the goal is to assign a class label to each pixel of the image. Semantic segmentation of the classified images with the model created above is its natural downstream application. In doing so, full utilization of the QSorter pipeline can be achieved, where along with the immediate results of seed embryo pixel identification, these results can be combined with hyperspectral imaging data in a simple regression problem to pair results in segmentation with results in direct imaging.

#### 2.4.1 Evaluation metrics

The Sørensen–Dice coefficient, also known as the Dice Similarity Coefficient (DSC), is a metric often used in segmentation tasks to evaluate the spatial overlap between two image masks [47]. It is given by the equation below:

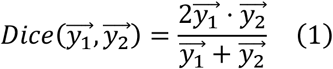

Here, 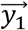 and 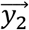 are the mask tensors flattened to one dimension. In statistical validation for computer vision tasks, DSC is often preferred over the pixel accuracy metric because DSC ignores true negatives, and pixel classes are often heavily biased toward the (negative) background, especially in binary semantic segmentation.

#### 2.4.2 Model details

U-Net is a convolutional neural network commonly used for semantic segmentation tasks [35×48]. It consists of a symmetric encoder-decoder pair, where the encoder down-samples while increasing the number of channels until a bottleneck tensor, from which the decoder up-samples while reducing the number of channels. For the segmentation task, we used U-Net with ResNet50 used as the encoder to both utilize and compare the self-supervised weights learned during the classification phase, as has been implemented in the literature to considerable advantage [36×49]. In this architecture, the encoder and decoder are not symmetric, as opposed to standard U-Net without a backbone, but skip connections are still fully implemented by limiting the depth of the encoder. **Figure 7** shows a U-Net with a ResNet50 as its encoder and four sets of multi-channel feature maps.

**Figure 7.**
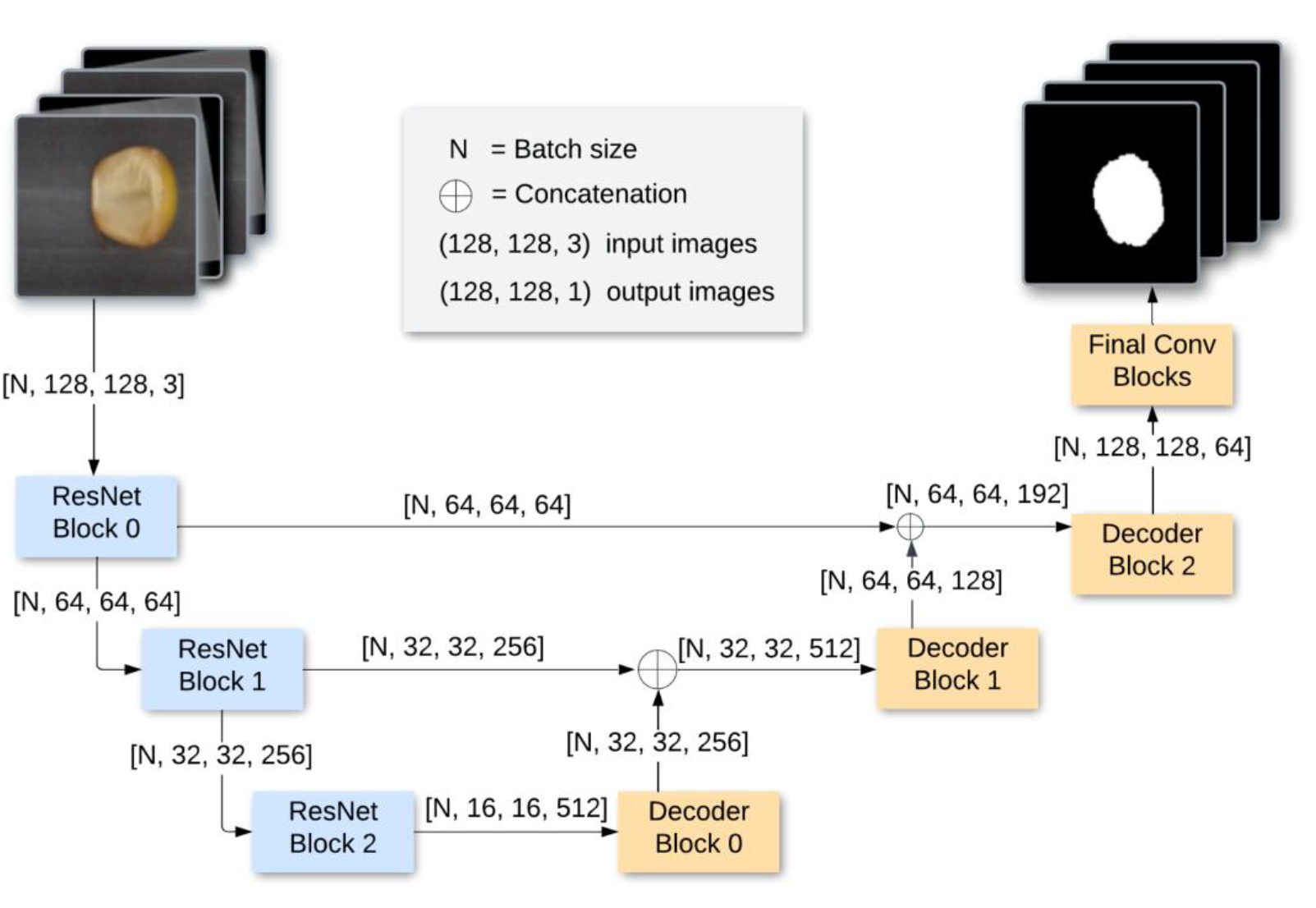
U-Net with ResNet50 backbone using filters of channel dimensions [64, 128, 256, 512]

#### 2.4.3 Data augmentation

Data augmentation was applied to each training batch to increase the set of distinct training images and to reduce overfitting. Augmentations were coupled between any RGB image and its mask. All augmentations were executed with a 50% application chance. These consisted of combinations of the following: 1) horizontal flip across the vertical middle axis, 2) paired brightness and contrast transform with an application factor uniformly selected from [-0.2, 0.2], and 3) paired scaling and shearing affine transform, the scaling factor uniformly selected from [0.75, 1] and the shear angle uniformly selected from [-π/6, π/6]. **Figure 8** shows an example of an augmented image-mask pair.

**Figure 8.**
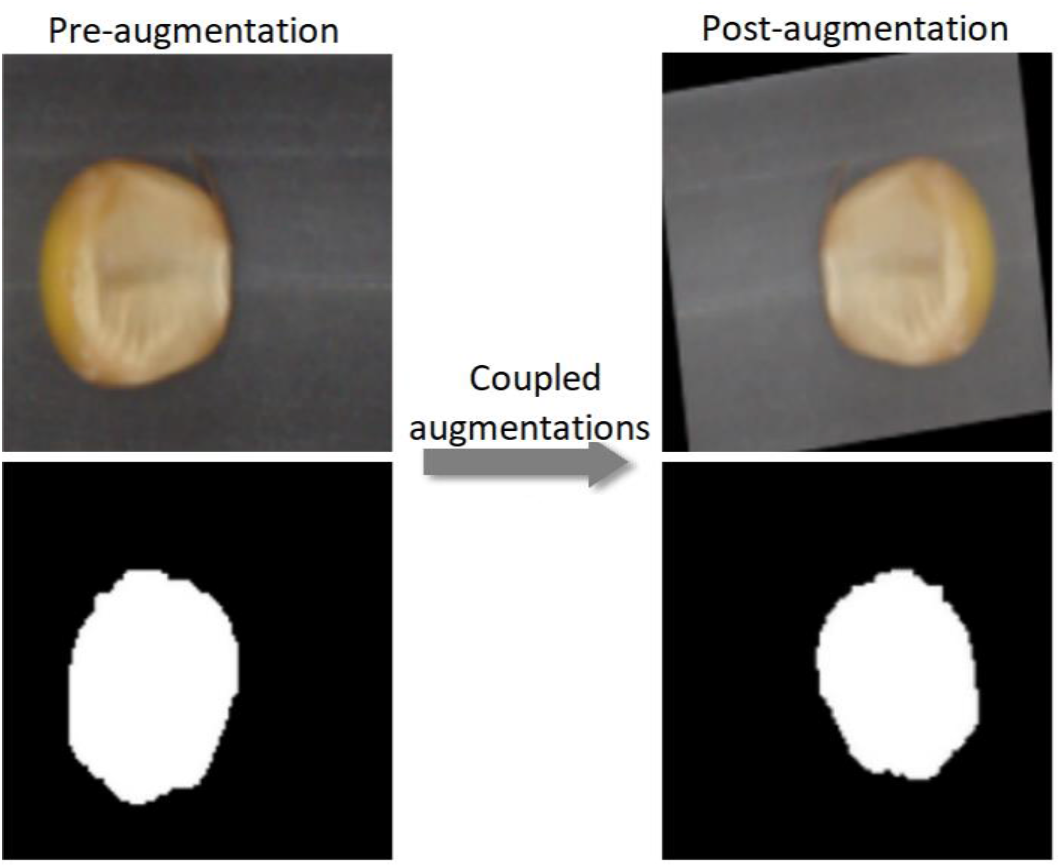
Coupled augmentations for an RGB image and its mask.

#### 2.4.4 Training process

Due to the smaller size of the segmentation dataset compared to the classification dataset, ten-fold cross-validation was performed. Using ten folds, ten models were created separately for each backbone and each set of hyperparameters, repeated for each of the three weight initialization types, each trained on a train/validation split of 288/32. With every ten folds, the highest average Dice score across all ten was collected. A model with this set of best-performing hyperparameters was trained on all training data without a validation set for 300 epochs. This model was then evaluated on the full test set. **Figure 9** illustrates the cross-validation process. Training and experiments were completed using Google Colab with NVIDIA Tesla T4 and K80 GPUs on 32 GB RAM.

**Figure 9.**
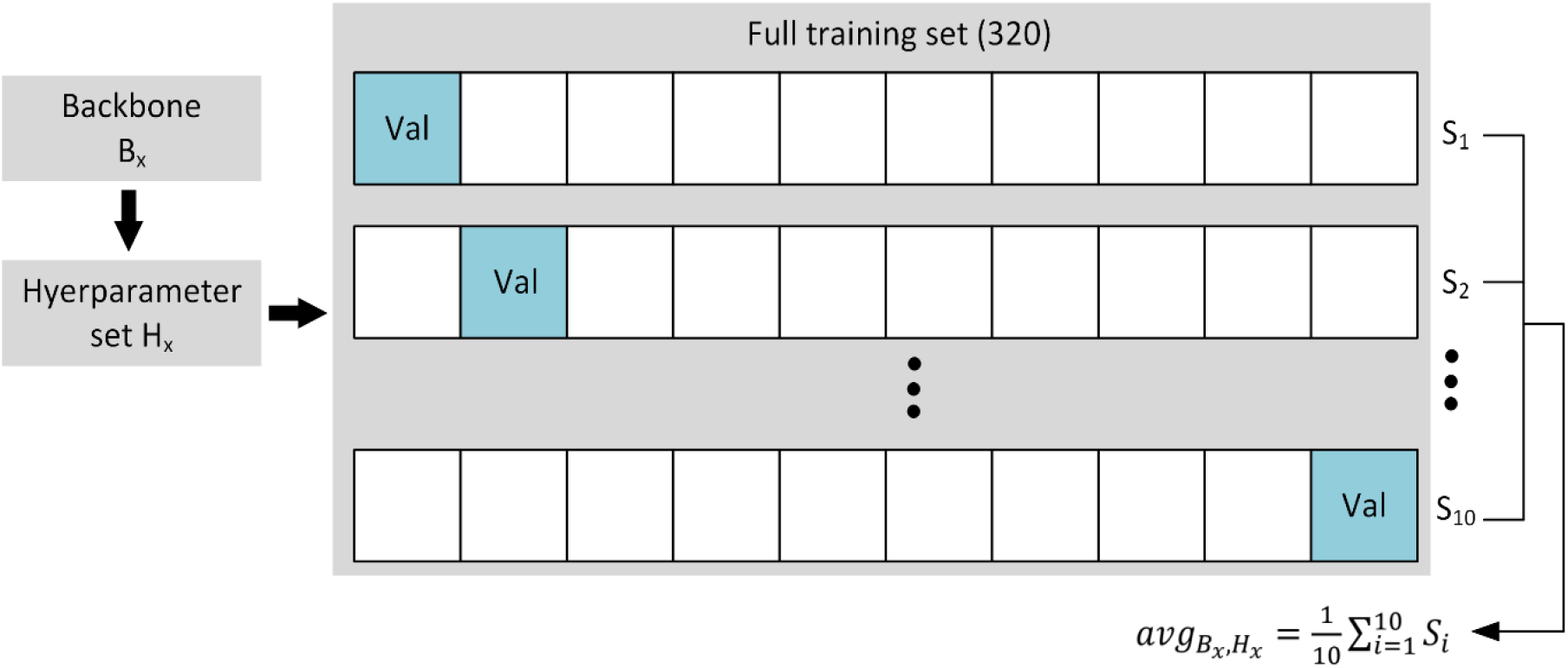
Cross-validation model selection procedure.

## 3. RESULTS AND DISCUSSION

### 3.1 Classification results

#### 3.1.1 Finetuning evaluation

After end-to-end finetuning, both SimCLR and NNCLR were more annotation-efficient and performed better than purely transfer learning-based methods, as shown in **Figure 10**. Listing test results from greatest to least utilization of total available annotated data, the NNCLR-pretrained model had accuracies of 85.6%, 83.8%, 81.6%, and 80.2%; the SimCLR-pretrained model had accuracies of 85.2%, 84.0%, 81.4%, and 81.8%; and the ImageNet-initialized model had accuracies of 84.0%, 81.4%, 77.2%, and 76.8%. At every annotation percentage, the self-supervised models outperformed all other models, with the largest difference at 1% annotation, where the SimCLR-pretrained model outperformed the ImageNet-based model by 5.0%. We remind the reader that the total available annotated data is only around 5% of the total data (2697 annotated images out of 44,286 total images). SSL pretraining provides a significant boost in model performance, especially at very low total annotated data availability; for instance, a 10% usage of annotated data represents just 270 annotated images!

**Figure 10.**
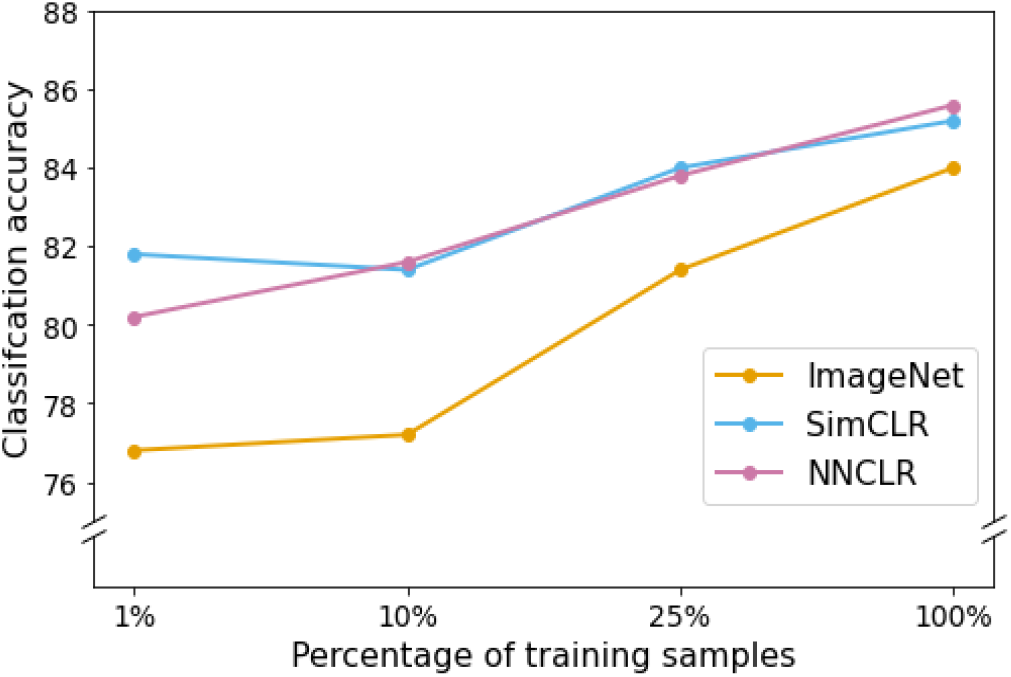
Classification accuracy versus percentage of training samples for three types of weight initializations.

#### 3.2 Comparisons

Models pretrained with contrastive SSL outperformed transfer learning models in every trial and between all data splits. **Table 1** shows the performances of each model compared to the ImageNet-pretrained model. The results of the SimCLR and NNCLR pretrained models outperforming the transfer learning model and being more annotation efficient are clear. The performances of NNCLR and SimCLR were similar to each other among the four annotation percentages, but in training on the full dataset, NNCLR performed slightly better, while SimCLR was more efficient at the lowest data split.

**Table 1.** Relative performance by the accuracy of SimCLR-pretrained and NNCLR-pretrained models as compared to ImageNet preloaded model. Each entry represents a performance gap in Fig. 9.

| Pretraining | Percentage annotated data used (available annotated dataset size is 2697) |  |  |  |
| --- | --- | --- | --- | --- |
|  | 1% | 10% | 25% | 100% |
| SimCLR | +3.4% | +4.2% | +2.6% | +1.2% |
| NNCLR | +5.0% | +4.4% | +2.4% | +1.4% |

### 3.2 Segmentation results

**Table 2** shows the test dataset evaluation results after the best models were selected and then finetuned, according to data from the previous three tables. It also shows the validation statistics and hyperparameter set for the chosen model. In **Appendix, Tables A1, A2**, and **A3** show the averaged results from 10-fold cross-validation on U-Net with a ResNet50 backbone from weights pretrained with SimCLR, pretrained with NNCLR, and pretrained from ImageNet, respectively. **Table A4** shows the selected models’ hyperparameter set. The U-Net with a SimCLR-pretrained backbone trained at 1e-04 LR and four encoder-decoder filters performed best, with a test DICE score of 0.81 compared to an ImageNet-pretrained backbone at 0.78 DICE score. The results from this section have a twofold implication: 1) they show U-Net with a backbone loaded with self-supervised pretrained weights can perform well, producing ∼0.81 Dice score, and 2) they show semantic segmentation with these backbones outperform those pre-trained with ImageNet. **Figure 11** displays three representative results from the segmentation model, including the predicted mask, the true mask, and the input RGB image.

**Table 2.**
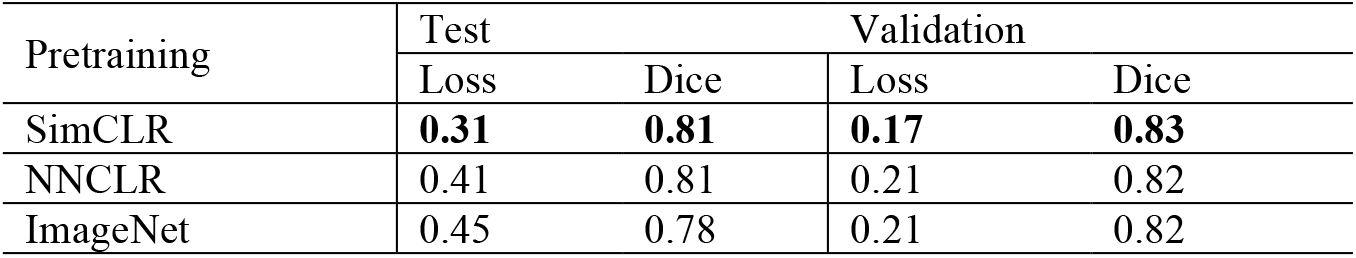
Loss and Dice scores for best hyperparameter sets for each weight initialization type. The validation column shows the average scores corresponding to each chosen hyperparameter set.

| Pretraining | Test |  | Validation |  |
| --- | --- | --- | --- | --- |
|  | Loss | Dice | Loss | Dice |
| SimCLR | **0.31** | **0.81** | **0.17** | **0.83** |
| NNCLR | 0.41 | 0.81 | 0.21 | 0.82 |
| ImageNet | 0.45 | 0.78 | 0.21 | 0.82 |

**Figure 11.**
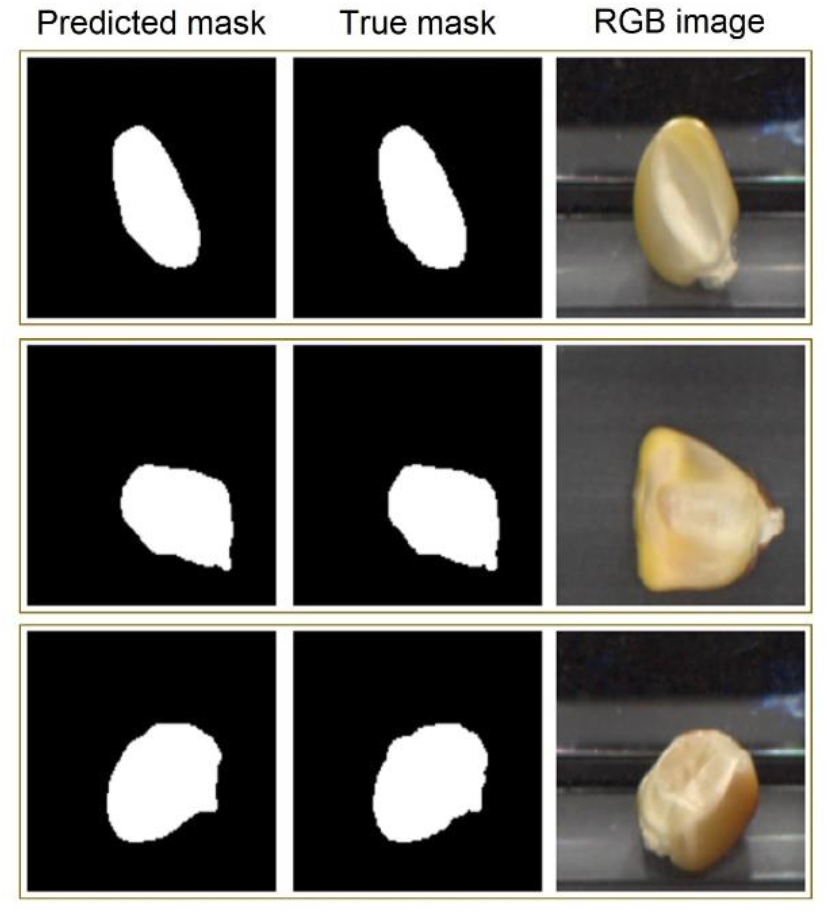
Three representative rows of segmentation inputs and outputs. The first column shows the predicted mask, the second shows the true mask, and the third shows the RGB input image.

## 4. CONCLUSION

From training contrastive learning models and comparing them with purely supervised and transfer learning methods, we found that self-supervised learning produces successful representations of an agricultural dataset applicable for downstream applications. We showed that NNCLR and SimCLR methods performed significantly better than their supervised counterparts, especially for the classification problem. These results also support the usage of strong augmentations in contrastive learning—far stronger than in end-to-end finetuning. In segmentation, self-supervised methods significantly improved over ImageNet pretraining, resulting in accurate masking capabilities and relative embryo size calculation. The combined results further show the transferable nature of self-supervised training. In particular, we illustrated that a single SSL-pretrained model (ResNet50 backbone) could be finetuned and used for two distinct downstream tasks – classification and segmentation. Furthermore, SSL pretraining allowed us to train models with very competitive performance even with very low amounts of total annotated data, for instance, with less than 1% (∼400 out of 44000 total images) of annotation. Thus, we have demonstrated that self-supervised learning provides a meaningful path forward in advancing agricultural efficiency with computer vision and machine learning.

## ACKNOWLEDGEMENT

This work was partially supported by AI Institute for Resilient Agriculture (USDA-NIFA #2021-67021-35329), COALESCE: COntext Aware LEarning for Sustainable CybEr-Agricultural Systems (CPS Frontier # 1954556). BG and TL acknowledge support from PSI faculty fellowship. We thank Dr. Candice Gardner at the USDA-ARS Plant Introduction Station.

## APPENDIX

**Table A1.** Average validation metrics on U-Net from SimCLR self-supervised pretrained weights

| Learning rate | 1e-03 |  |  | 1e-04 |  |  | 1e-05 |  |  |
| --- | --- | --- | --- | --- | --- | --- | --- | --- | --- |
| Channel count | 3 | 4 | 5 | 3 | 4 | 5 | 3 | 4 | 5 |
| BCE Loss | 0.132 | 0.163 | 0.206 | 0.135 | 0.171 | 0.213 | 0.099 | 0.116 | 0.155 |
| Accuracy | 0.963 | 0.965 | 0.964 | 0.962 | 0.967 | 0.966 | 0.961 | 0.963 | 0.963 |
| Dice Score | 0.797 | 0.807 | 0.810 | 0.761 | 0.828 | 0.821 | 0.764 | 0.797 | 0.771 |

**Table A2.** Average validation metrics on U-Net from NNCLR self-supervised pretrained weights

| Learning rate | 1e-03 |  |  | 1e-04 |  |  | 1e-05 |  |  |
| --- | --- | --- | --- | --- | --- | --- | --- | --- | --- |
| Channel count | 3 | 4 | 5 | 3 | 4 | 5 | 3 | 4 | 5 |
| BCE Loss | 0.131 | 0.164 | 0.163 | 0.163 | 0.216 | 0.181 | 0.100 | 0.128 | 0.173 |
| Accuracy | 0.950 | 0.954 | 0.972 | 0.961 | 0.967 | 0.966 | 0.954 | 0.971 | 0.964 |
| Dice Score | 0.792 | 0.791 | 0.794 | 0.793 | 0.824 | 0.820 | 0.723 | 0.800 | 0.802 |

**Table A3.** Average validation metrics on U-Net from ImageNet pretrained weights

| Learning rate | 1e-03 |  |  | 1e-04 |  |  | 1e-05 |  |  |
| --- | --- | --- | --- | --- | --- | --- | --- | --- | --- |
| Channel count | 3 | 4 | 5 | 3 | 4 | 5 | 3 | 4 | 5 |
| BCE Loss | 0.194 | 0.200 | 0.202 | 0.200 | 0.208 | 0.183 | 0.103 | 0.196 | 0.182 |
| Accuracy | 0.965 | 0.967 | 0.966 | 0.959 | 0.965 | 0.966 | 0.960 | 0.964 | 0.956 |
| Dice Score | 0.810 | 0.824 | 0.826 | 0.783 | 0.819 | 0.798 | 0.765 | 0.676 | 0.771 |

**Table A4.** Hyperparameter values for best segmentation models for each weight initialization type as found by cross-validation

|  | Filters | Learning rate |
| --- | --- | --- |
| SimCLR | 4 | 1e-04 |
| NNCLR | 4 | 1e-04 |
| ImageNet | 5 | 1e-03 |

